# A protocol for good quality genomic DNA isolation from formalin-fixed paraffin-embedded tissues without using commercial kits

**DOI:** 10.1101/2021.07.23.452892

**Authors:** Fazlur Rahman Talukdar, Irena Abramović, Cyrille Cuenin, Christine Carreira, Nitin Gangane, Nino Sincic, Zdenko Herceg

## Abstract

DNA isolation from formalin-fixed paraffin-embedded (FFPE) tissues for molecular analysis has become a frequent procedure in cancer research. However, the yield or quality of the isolated DNA is often compromised, and commercial kits are used to overcome this to some extent. We developed a new protocol (IARCp) to improve better quality and yield of DNA from FFPE tissues without using any commercial kit. To evaluate the IARCp’s performance, we compared the quality and yield of DNA with two commercial kits, namely NucleoSpin® DNA FFPE XS (MN) and QIAamp DNA Micro (QG) isolation kit. Total DNA yield for QG ranged from 120.0 – 282.0 ng (mean 216.5 ng), for MN: 213.6 – 394.2 ng (mean 319.1 ng), and with IARCp the yield was much higher ranging from 775.5 – 1896.9 ng (mean 1517.8 ng). Moreover, IARCp has also performed well in qualitative assessments. Overall, IARCp represents a novel approach to DNA isolation from FFPE which results in good quality and significant amounts of DNA suitable for many downstream genome-wide and targeted molecular analyses. Our proposed protocol does not require the use of any commercial kits for isolating DNA from FFPE tissues, making it suitable to implement in low-resource settings such as low and middle-income countries (LMICs).

## 1. INTRODUCTION

Formalin-fixed paraffin-embedded (FFPE) tissue samples represent a preserved specimen routinely used for cancer diagnosis and research. The process of dehydrating the tissue and fixating it with formalin, and then embedding the tissue in paraffin wax facilitates the cutting of thin tissue sections for precise histomorphological, immunohistochemical, and other clinical analyses ^1^. An archive of FFPE tissues around the world is a valuable source for research, especially in the cancer field, enabling remarkable advances in understanding tumor biology and molecular biomarkers ^2^. The routine clinical workflows in hospitals do not typically have the facility to collect clinically relevant fresh tissues, which usually yield optimum quality and quantity of nucleic acids for research purposes. Therefore, the use of FFPE tissues to perform various molecular analyses becomes a critical necessity^1^. Using FFPE for genetic analysis is one of the most-used applications in cancer research, especially since next-generation sequencing (NGS) is finding its place in the clinic. However, there are several technical challenges with DNA extraction from FFPE samples that are affecting the downstream genomic analysis. Apart from preanalytical factors in FFPE preparation affecting DNA analysis ^3^, partial DNA degradation and DNA binding to amino acids occur in the light of formalin fixation ^2,4^. Therefore, DNA recovered from FFPE is significantly fragmented, both due to formalin fixation and prolonged storage, which impairs polymerase chain reaction (PCR) and NGS performance, as well as does the potential contamination with inhibitors ^5,6^. Moreover, different extraction methods lead to variable DNA quantity and quality which may influence the results ^7,8^. The DNA yield is of special importance when isolating from minute amounts of tissue such as biopsy samples, which are in their nature very small, while also limited in the amount which is available for isolation since multiple analyses are performed on the same FFPE sample ^8^.

Numerous commercial kits for DNA isolation from FFPE, both manual and automatic, are available on the market, as well as in-house protocols, with different performance characteristics ^2,9^. Also, there is no lack of comparison in the literature, with substantial differences observed in the terms of DNA quality and quantity ^8,10–14^. However, the cancer research field still faces a challenge to establish a robust, reliable, and reproducible isolation protocol for obtaining enough good-quality DNA from minimal amounts of FFPE tissue, especially for downstream analyses. Moreover, there is a lack of cost-effective techniques (without using commercial kits) for FFPE tissue DNA isolation.

To address this challenge, we developed at the International Agency for Research on Cancer a protocol (IARCp) that offers remarkably good DNA recovery, purity, and amplifiability suitable for a broad range of downstream analyses. The present experimental plan aimed to compare IARCp with two commercial protocols Macherey-Nagel’s NucleoSpin® DNA FFPE XS kit (MN), and Qiagen’s QIAamp DNA Micro isolation kit (QG), for DNA isolation from FFPE tissues.

## 2. MATERIALS AND METHODS

### 2.1 Samples

A total of six FFPE tissue blocks of the esophageal biopsy included in this experiment was collected from Mahatma Gandhi Institute of Medical Sciences (MGIMS), India for a multi-center study on esophageal cancer^15^. Ethical approval for the study was granted by the MGIMS local ethical committee and IARC Ethical Committee approved this project under approval number 16-25.

### 2.2 DNA extraction

From each FFPE sample, eleven 10 µm thick sections were cut and placed on a glass slide. The first and the last cut sections were stained with hematoxylin-eosin for the routine pathological examination. The remaining nine intermediate sections were alternatively collected on Superfrost® Plus & Colorfrost® (ROTH SOCHIEL EURL) adhesion slides for the isolation with three different protocols, as shown in **Figure 1**. Three alternate tissue sections were collected for the three different protocols (as shown in Figure 1) to avoid any possible bias related to the tissue surface area present in each section used for the isolation. The use of SuperFrost Plus adhesion slides is essential here to ensure the proper attachment of tissue sections with the slides through different deparaffinization and rehydration steps. DNA was isolated from cuts according to the manufacturer’s protocol, for NucleoSpin® DNA FFPE XS (Macherey-Nagel) and QIAamp DNA Micro (Qiagen) isolation kit, and according to our protocol, as shown in **Figure 1** and **Table 1**. Deparaffinization was done by submerging the slides to Xylene, 100 ethanol (EtOH), 95% EtOH, 70% EtOH and deionized water for 5 minutes each.

**Table 1.** Protocol for DNA isolation from FFPE developed at IARC (IARCp).

| Steps | Description |
| --- | --- |
| **(1)** Rehydration | Deparaffinized slides with tissue sections were kept submerged for 48 hours in PBS or TBS buffer (pH=7.2) at 4°C |
| **(2)** Tissue homogenization | The tissue was scraped and homogenized in small pieces using a sterile scalpel and transferred into a tube with 500 µL of TES buffer |
| **(3)** Tissue digestion | 20 µl proteinase K (10 mg/ml) was added to the tubes and incubated at 56°C in a hot plate/water bath until the tissue is dissolved completely.<br>(*If the tissue is not properly dissolved after 2-3 hours of incubation, then another 10 µl proteinase K can be added and incubated overnight until the tissue is completely dissolved) |
| **(4)** Vortex | After dissolving the tissue, the tubes were vortexed shortly and spin down the content |
| **(5)** Adding concentrated salt (6M NaCl) | Then 200 µL of 6M NaCl was added to the tubes and vortexed for 5 minutes |
| **(6)** Centrifuge; (≥13000 rpm) | The tubes were then centrifuged at full speed (≥13000 rpm) for 10 min |
| **(7)** Supernatant collected | The supernatant (around 700 µL) was transferred into a clean tube without disturbing the pellet |
| **(8)** Adding isopropanol/ DNA precipitation | 500 µL of isopropanol was added to the supernatant |
| **(9)** Centrifuge; (≥13000 rpm) to get DNA pellet | The contents were vortexed for 2 minutes and centrifuged at full speed (≥13000 rpm) for 15 min to get DNA pellet |
| **(10)** 70% ethanol wash | The supernatant was decanted and 500 µL of 70% ethanol (EtOH) was added to wash the DNA pellet by tapping the bottom of the tube several times |
| **(11)** Centrifuge; (≥13000 rpm) | The tubes were centrifuged at full speed (≥13000 rpm) for 15 min |
| **(12)** Carefully discarding ethanol | The EtOH was discarded by inverting the tube (carefully*) and the pellet was dried in a sterile environment until EtOH has evaporated.<br>(*This is a very critical step to be very careful as the pellet remains loosely bound to the tube. This step needs extra care while discarding the EtOH to avoid losing pellet with EtOH) |
| **(13)** DNA dissolved in TE buffer | 30 µL of nuclease-free water or TE (Tris-EDTA) buffer was added and gently mixed the DNA with the solution |
| **(14)** DNA quantification & storage | The isolated DNA was quantified and stored at a -20°C or -80°C freezer before further use. |

**Figure 1.**
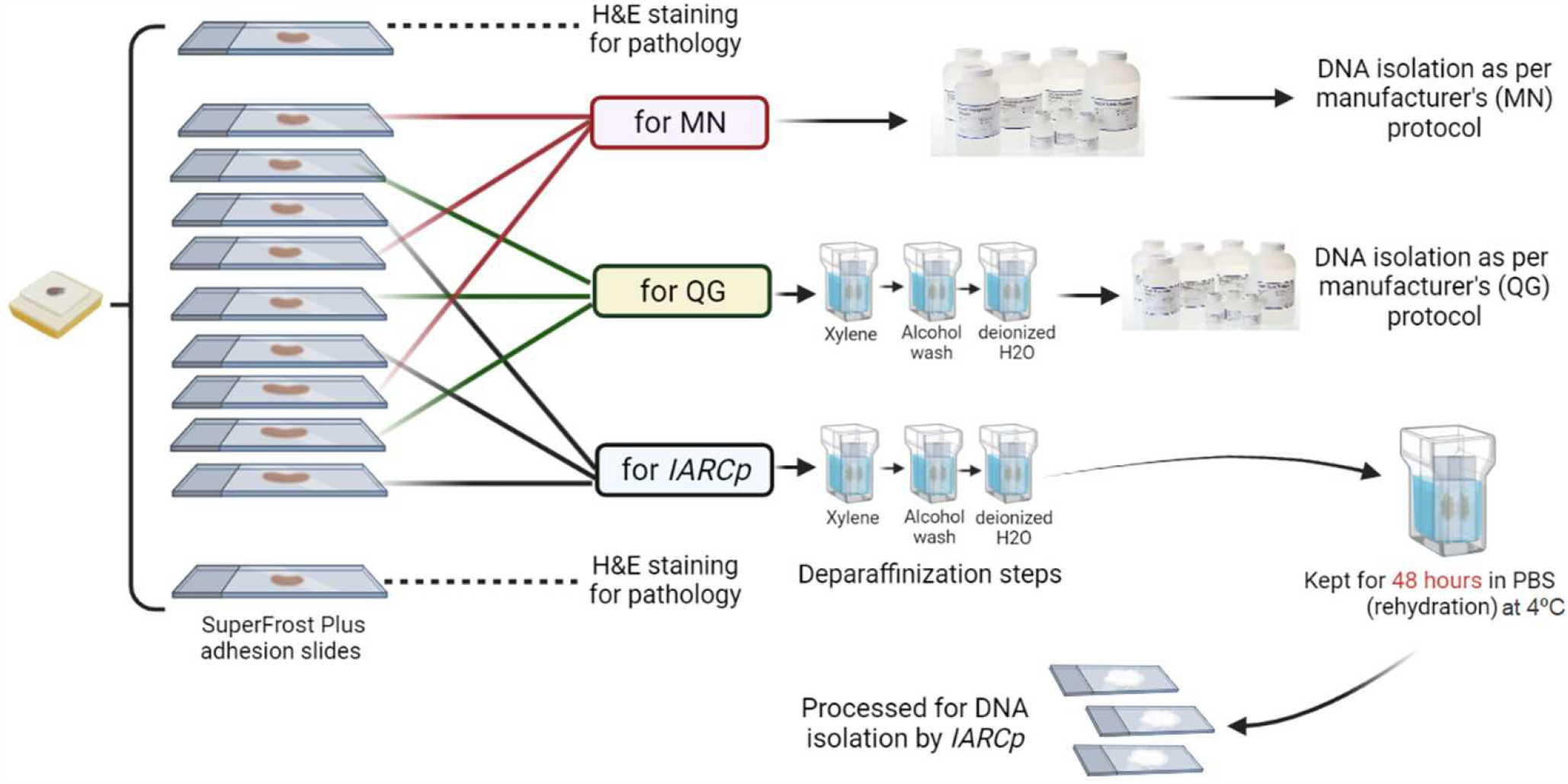
Study design - Three alternate 10 µm paraffin sections were taken for each DNA isolation protocol as shown in the figure. [**MN**: Macherey-Nagel (NucleoSpin® DNA FFPE XS); **QG**= Qiagen (QIAamp DNA Micro) and **IARCp**: International Agency for Research on Cancer developed Protocol]

After deparaffinization, samples were left to rehydrate by submerging them in a buffer (phosphate or Tris; pH=7.2) for 48 hours at 4°C. After the rehydration step, the tissue sections were scrapped from slides and homogenized. The homogenized tissues were transferred to a tube containing 500 µL TES buffer (50mM Tris-HCl pH=8; 100mM EDTA; 100mM NaCl; 1% SDS) and samples were left to solubilize at 56°C with proteinase K. Protein precipitation was performed with saturated 6M NaCl and DNA was precipitated with EtOH. Elution volumes were 30 µL in all protocols. The step-by-step DNA isolation by IARCp is described in detail in **Table 1**.

### 2.3 DNA quantification and quality control

All DNA samples were quantified by fluorometry using Qubit ™ dsDNA HS Assay Kit (Life Technologies, Carlsbad, California, US) on Qubit™ 3 Fluorometer (Invitrogen, Life Technologies) as per the manufacturer’s instructions, and assessed for purity by NanoDrop 8000 Spectrophotometer (Thermo Scientific) 260/280 absorbance ratio measurements, in triplicate.

Apart from the Qubit assay, for the quality control analysis of DNA, Infinium HD FFPE QC Assay (Illumina, Inc.) was used by performing a quantitative PCR of FFPE DNA on CFX96™ Real-Time PCR Detection System (BioRad). Subsequent data analysis was performed as per the manufacturer’s instructions. The ΔCq was calculated to evaluate the quality of isolated DNA, since values ΔCq > 5 are not suitable for further downstream processing for Infinium HD FFPE Restore Protocol (Illumina, Inc.) and Infinium MethylationEPIC array (Illumina, Inc.). A value ΔCq < 5 ensures the better quality of isolated DNA that is suitable for various targeted and genome-wide analyses.

### 2.4 Statistical analysis

The differences in total DNA yield between isolation protocols were analyzed with a one-way analysis of variance (ANOVA) and Student’s t-test for pairwise difference in R studio (version 4.0.5). The p values < 0.05 were considered statistically significant.

## 3. RESULTS

### 3.1 DNA yield

All three isolation methods recovered measurable amounts of DNA from the six tissue samples used in the experiment. The DNA quantifications were done by the Qubit fluorometer as per the manufacturer’s instruction. We observed variable results among the three protocols, where IARCp showed the best performance in comparison with QG and MN, which gave a significantly lower yield. Total DNA yield for QG ranged from 120.0 – 282.0 ng (mean 216.5 ng), for MN: 213.6 – 394.2 ng (mean 319.1 ng), and with IARCp it was much higher ranging from 775.5 – 1896.9 ng (mean 1517.8 ng) (**Figure 2A** and **Supplementary Table S1**). DNA isolated with IARCp was significantly more abundant than the one isolated with QG or MN, with a p-value < 0.0001.

**Figure 2.**
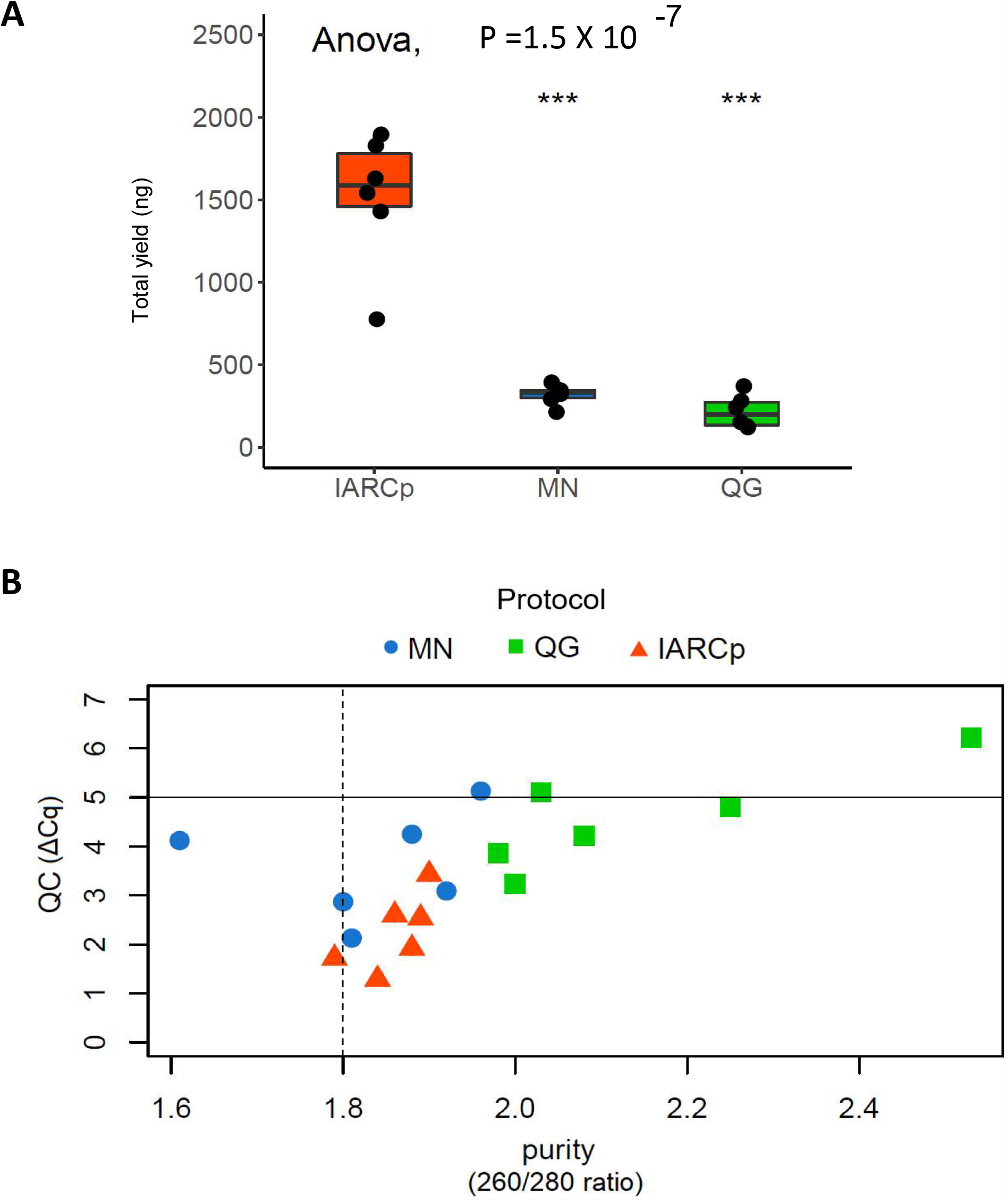
DNA yield and purity; **A)** Total DNA yield in nanograms (ng) with the three protocols; **B)** Volcano plot showing quality control step with Δ Ct values on Y-axis and the protocol tested in X-axis. The line in X-axis at 1.8 is ideal for DNA purity and the line in Y-axis at Δ Ct≥ 5 is the threshold for poor quality DNA. [**MN**: Macherey-Nagel (NucleoSpin® DNA FFPE XS); **QG**= Qiagen (QIAamp DNA Micro) and **IARCp**: International Agency for Research on Cancer developed protocol] *** P< 0.0001; Student’s t-test

### 3.2 DNA quality

The ratio of absorbance of DNA at 260/280 nm wavelength is usually used to estimate the purity of DNA and a value ∼1.8 is considered ideal for DNA purity ^16^. Among the three protocols tested, IARCp also exhibited better performance in the terms of DNA quality as the average absorbance ratio at 260/280 nm for all six samples was 1.86, MN protocol also exhibited a similar average value of 1.83. However, the average ratio for the QG protocol was 2.15, which is not an ideal value for DNA (**Figure 2B**). Additionally, we also performed Infinium HD FFPE QC Assay to evaluate the extent of DNA degradation in the isolates. All samples isolated with IARCp showed ΔCq < 5, suggesting lesser degradation of the isolated DNA. Moreover, five out of six samples isolated with IARCp exhibited ΔCq <3, further signifying the better quality of the isolated DNA. Regarding the protocol MN, five out of six samples showed ΔCq < 5 and one sample had ΔCq > 5 (poor quality). Among the samples isolated with QG protocol, four samples showed ΔCq < 5, and two of them were of poor quality (ΔCq > 5). These results are presented in **Figure 2B** and **Supplementary Table S1**.

## 4. DISCUSSION

DNA isolation from FFPE blocks has become a routine practice in cancer research but is often compromised in the terms of yield or quality. Moreover, it is difficult to isolate DNA from FFPE tissues without using commercial kits for molecular analyses. To address this, we developed a new protocol (IARCp) to improve better quality and yield of DNA without using any commercial kit. To evaluate the IARCp’s performance, we compared it with two commercial kits – Qiagen’s tissue DNA isolation protocol (protocol QG) and Macherey-Nagel’s FFPE DNA isolation protocol (protocol MN). IARCp showed exceptionally better performance as regards DNA quantity, with three times as much DNA isolated from the same specimen. Moreover, DNA purity, assigned as absorbance ratio 260/280 nm, was much better for IARCp than QG, and more consistent for IARCp than MN. As for DNA passing the quality control test, both IARCp and MN showed very good performance, while QG did not satisfy.

MN and QG use silica columns for DNA isolation and purification due to which they require short performance time which is their advantage. Besides, MN avoids the use of harsh chemicals for deparaffinization. QG is not specifically designed for DNA isolation from FFPE but was developed for very small tissue samples like the one used in this study. Not so optimal results of QG could be ascribed to the fact that it was not designed for FFPE, even though it has been shown that Qiagen’s kits not originally intended for DNA isolation from FFPE could be used for this purpose ^17^. However, the expected DNA yield when using Qiagen’s kit for FFPE is still quite low compared to IARCp, also when considering the amount of tissue involved ^10,11,18^. Although MN is being used among researchers, the data on its comparison with other kits and protocols is rather scarce ^2,19^.

The critical difference from commercial kits is that IARCp requires an additional rehydration step where samples are submerged in a buffer for 48 hours after the deparaffinization. This could be a limitation of this protocol due to the requirement of additional time, yet this step allows proper rehydration of the tissue. Additional tissue rehydration steps could result in better tissue or cell lysis, leading to the proper release of nucleic acids from the tissues in the lysis solution. This is probably the reason why IARCp yields such significantly abundant and good-quality DNA by increasing water content while reducing stiffness within the tissue section ^20^. Also, IARCp outperforms other in-house developed protocols suggested for FFPE isolation, based on DNA yield ^13,14^. In the future, it would be useful to apply IARCp for different FFPE tissue types for comparison, but for now, we have had a great experience when using it from a whole range of cancer and healthy tissues (data not shown).

In conclusion, we compared the analytical performance of our developed IARCp with two commercially available kit protocols for DNA isolation from FFPE tissues. IARCp has several advantages such as the following. Firstly, it has shown exceptionally better performance in terms of DNA quantity and quality, while also being appropriate for very small amounts of tissue. Secondly, in IARCp we do not use phenol and chloroform-based DNA extraction which are highly toxic reagents, making the protocol safer for the experiment performer. Finally, our protocol does not require the use of any commercial kits which makes it suitable to implement in low-resource settings such as low and middle-income countries (LMICs). Therefore, IARCp represents a novel approach to DNA isolation from FFPE which results in good quality significant amounts of DNA suitable for many downstream analyses. The remarkable performance of IARCp in terms of DNA quantity and quality enables the use of minuscule FFPE tissue amounts to be used for various downstream analyses from whole-exome sequencing to target mutation detection using sequencing analyses. Moreover, we have used IARCp derived DNA for DNA methylation analyses such as methylome analysis with Infinium MethylationEPIC array and targeted DNA methylation analysis using pyrosequencing^15^.

## IARC DISCLAIMER

Where authors are identified as personnel of the International Agency for Research on Cancer / World Health Organization, the authors alone are responsible for the views expressed in this article and they do not necessarily represent the decisions, policy or views of the International Agency for Research on Cancer / World Health Organization.

## ACKNOWLEDGEMENTS

The work reported in this article was undertaken by F.R. Talukdar partly during the tenure of a Postdoctoral Fellowship from the International Agency for Research on Cancer (IARC), partially supported by the EC FP7 Marie Curie Actions—People— Co-funding of regional, national, and international programs (COFUND).

## REFERENCES

1. Mathieson, W. & Thomas, G. Using FFPE Tissue in Genomic Analyses: Advantages, Disadvantages and the Role of Biospecimen Science. Current Pathobiology Reports (2019) doi:10.1007/s40139-019-00194-6.

2. Patel, P. G. et al. Reliability and performance of commercial RNA and DNA extraction kits for FFPE tissue cores. PLoS ONE (2017) doi:10.1371/journal.pone.0179732.

3. Bass, B. P., Engel, K. B., Greytak, S. R. & Moore, H. M. A review of preanalytical factors affecting molecular, protein, and morphological analysis of Formalin-Fixed, Paraffin-Embedded (FFPE) tissue: How well do you know your FFPE specimen? Archives of Pathology and Laboratory Medicine (2014) doi:10.5858/arpa.2013-0691-RA.

4. Angelo Fortunato, Diego Mallo, Shawn M Rupp, Lorraine M King, Timothy Hardman, Joseph Y Lo, Allison Hall, Jeffrey R Marks, E Shelley Hwang, C. C. M. A new method to accurately identify single nucleotide variants using small FFPE breast samples. Brief. Bioinform. (2021).

5. Dietrich, D. et al. Improved PCR Performance Using Template DNA from Formalin-Fixed and Paraffin-Embedded Tissues by Overcoming PCR Inhibition. PLoS ONE (2013) doi:10.1371/journal.pone.0077771.

6. Watanabe, M. et al. Estimation of age-related DNA degradation from formalin-fixed and paraffin-embedded tissue according to the extraction methods. Exp. Ther. Med. (2017) doi:10.3892/etm.2017.4797.

7. Lu, X. J. D., Liu, K. Y. P., Zhu, Y. S., Cui, C. & Poh, C. F. Using ddPCR to assess the DNA yield of FFPE samples. Biomol. Detect. Quantif. (2018) doi:10.1016/j.bdq.2018.10.001.

8. Kresse, S. H. et al. Evaluation of commercial DNA and RNA extraction methods for high-throughput sequencing of FFPE samples. PLoS ONE (2018) doi:10.1371/journal.pone.0197456.

9. Kocjan, B. J., Hošnjak, L. & Poljak, M. Commercially available kits for manual and automatic extraction of nucleic acids from formalin-fixed, paraffin-embedded (FFPE) tissues. Acta Dermatovenerol. Alp. Pannonica Adriat. (2015) doi:10.15570/actaapa.2015.12.

10. Mathieson, W., Guljar, N., Sanchez, I., Sroya, M. & Thomas, G. A. Extracting DNA from FFPE Tissue Biospecimens Using User-Friendly Automated Technology: Is There an Impact on Yield or Quality? Biopreservation Biobanking (2018) doi:10.1089/bio.2018.0009.

11. Sarnecka, A. K. et al. DNA extraction from FFPE tissue samples – a comparison of three procedures. Wspolczesna Onkol. (2019) doi:10.5114/wo.2019.83875.

12. Mcdonough, S. J. et al. Use of FFPE-derived DNA in next generation sequencing: DNA extraction methods. PLoS ONE (2019) doi:10.1371/journal.pone.0211400.

13. Okello, J. B. A. et al. Comparison of methods in the recovery of nucleic acids from archival formalin-fixed paraffin-embedded autopsy tissues. Anal. Biochem. (2010) doi:10.1016/j.ab.2010.01.014.

14. Ludyga, N. et al. Nucleic acids from long-term preserved FFPE tissues are suitable for downstream analyses. Virchows Arch. (2012) doi:10.1007/s00428-011-1184-9.

15. Talukdar, F. R. et al. Genome-Wide DNA Methylation Profiling of Esophageal Squamous Cell Carcinoma from Global High-Incidence Regions Identifies Crucial Genes and Potential Cancer Markers. Cancer Res. 81, 2612–2624 (2021).

16. Lucena-Aguilar, G. et al. DNA Source Selection for Downstream Applications Based on DNA Quality Indicators Analysis. in Biopreservation and Biobanking (2016). doi:10.1089/bio.2015.0064.

17. Bhagwate, A. V. et al. Bioinformatics and DNA-extraction strategies to reliably detect genetic variants from FFPE breast tissue samples. BMC Genomics (2019) doi:10.1186/s12864-019-6056-8.

18. Kalmár, A. et al. Comparison of Automated and Manual DNA Isolation Methods for DNA Methylation Analysis of Biopsy, Fresh Frozen, and Formalin-Fixed, Paraffin-Embedded Colorectal Cancer Samples. J. Lab. Autom. (2015) doi:10.1177/2211068214565903.

19. Adams, A. J. et al. DNA extraction method affects the detection of a fungal pathogen in formalin-fixed specimens using qPCR. PLoS ONE (2015) doi:10.1371/journal.pone.0135389.

20. Safa, B. N., Meadows, K. D., Szczesny, S. E. & Elliott, D. M. Exposure to buffer solution alters tendon hydration and mechanics. J. Biomech. (2017) doi:10.1016/j.jbiomech.2017.06.045.

