## Supplemental Table S1 for "A protocol for good quality genomic DNA isolation from formalin-fixed paraffin-embedded tissues without using commercial kits"

**Supplementary table S1**. Compiled results of DNA yield, purity, and quality

| Sample ID | Protocol used | Nanodrop 260/280 | Qubit (ng/µl) | total yield (ng) | Illumina QC (ΔCq) |
|---|---|---|---|---|---|
| **QG1** | QG | 2.25 | 8.08 | 242.4 | 4.81 |
| **QG2** | QG | 2.08 | 12.4 | 372 | 4.22 |
| **QG3** | QG | 2.53 | 4 | 120 | 6.22 |
| **QG4** | QG | 2.03 | 5.12 | 153.6 | 5.11 |
| **QG5** | QG | 1.98 | 4.3 | 129 | 3.86 |
| **QG6** | QG | 2 | 9.4 | 282 | 3.24 |
| **MN1** | MN | 1.96 | 11.6 | 348 | 5.13 |
| **MN2** | MN | 1.8 | 10.8 | 324 | 2.87 |
| **MN3** | MN | 1.61 | 13.14 | 394.2 | 4.12 |
| **MN4** | MN | 1.92 | 9.7 | 291 | 3.09 |
| **MN5** | MN | 1.88 | 7.12 | 213.6 | 4.25 |
| **MN6** | MN | 1.81 | 11.45 | 343.5 | 2.13 |
| **IARCp1** | IARCp | 1.86 | 54.34 | 1630.2 | 2.6 |
| **IARCp2** | IARCp | 1.89 | 61 | 1830 | 2.54 |
| **IARCp3** | IARCp | 1.79 | 51.44 | 1543.2 | 1.72 |
| **IARCp4** | IARCp | 1.88 | 63.23 | 1896.9 | 1.92 |
| **IARCp5** | IARCp | 1.9 | 25.85 | 775.5 | 3.43 |
| **IARCp6** | IARCp | 1.84 | 47.7 | 1431 | 1.29 |
